# Mapping Anhedonia-Related Damage Network: Insights for TMS Treatment

**DOI:** 10.1101/2025.02.11.637479

**Authors:** Chengfeng Chen, Shiying Wang, Yuan Liu, Xuezhen Zhang, Xinyan Rong, Jiang Wang, Jijun Wang, Ling Sun, Bin Zhang

**Author notes:** Equally contributed to this study as a co-first author (Chengfeng Chen, Shiying Wang, Yuan Liu). Corresponding author: Bin Zhang, Institute of Mental Health, Tianjin Anding Hospital, Tianjin Medical University, Tianjin, China,; Ling Sun, Institute of Mental Health, Tianjin Anding Hospital, Tianjin Medical University, Tianjin, China.

## Abstract

**Background:** Many studies have explored anhedonia-related functional connectivity (FC), but the findings remain inconsistent. There is a gap in identifying a consistent anhedonia-related damage network and applying it to TMS treatment.

**Methods:** We systematically reviewed studies on anhedonia-related functional connectivity and identified anhedonia-related brain damage locations. Using a novel functional connectivity network mapping approach applied to a large normative connectome dataset, we mapped these damage locations to anhedonia-related damage networks. Subsequently, transcriptomic analysis was conducted to uncover underlying molecular mechanisms. Additionally, we investigated the application of the anhedonia-related damage network in transcranial magnetic stimulation (TMS) treatment, focusing on changes in FC within this network following TMS treatment and its association with anhedonia improvement, as well as predicting TMS treatment efficacy based on baseline FC within the anhedonia-related damage network.

**Results:** A total of eight experiments from seven studies using the nucleus accumbens (NAc) as the seed were eligible for functional connectivity network mapping analysis. This study identified an anhedonia-related damage network, primarily characterized by disrupted functional connectivity between the NAc and the default mode network. Transcriptomic analysis revealed gene enrichment associated with synaptic signaling, neuronal development, ion transport, and actin cytoskeleton regulation. TMS treatment increased NAc functional connectivity within the anhedonia-related damage network in the response group, with these changes correlating with improvements in anhedonia. Furthermore, baseline NAc FC within this network demonstrated predictive potential for TMS treatment efficacy.

**Conclusion:** The anhedonia-related damage network was identified, emphasizing its underlying mechanisms and predictive value for TMS in treating anhedonia.

## 1. Introduction

Anhedonia, characterized by a profound loss of interest in or the inability to experience pleasure, often persists despite traditional antidepressant treatments (1, 2). It is also linked to poor responses to these treatments (3, 4), highlighting the need to target anhedonia-related neural pathways in the development of more effective therapeutic approaches.

Drawing on the contemporary understanding of depression as a disorder of brain circuits (5), functional connectivity (FC) has been recognized as crucial for treating anhedonia. Several studies have explored FC patterns associated with anhedonia. One study reported a negative correlation between FC between the nucleus accumbens (NAc) and the orbitofrontal cortex and the severity of anhedonia (6). Another study identified that greater anhedonia severity was linked to reduced FC between the NAc and a region extending from the caudate to the thalamus (7). Additionally, FC between the NAc shell-like subdivision and the subgenual anterior cingulate cortex or pregenual anterior cingulate cortex has also been reported to be negatively correlated with anhedonia severity (8). While these findings highlight important neural substrates, the neurobiological mechanisms underlying anhedonia remain inconsistent across studies, suggesting the need for further investigation.

Functional connectivity network mapping (FCNM) (9) is a novel approach to solving the heterogeneity of neuroimaging results. FCNM employs damage coordinates to map brain networks associated with specific symptoms, drawing on a large cohort of resting-state normative functional connectivity data (9). This method has been applied to identify brain structural and functional damage network localizations in suicide (10, 11), convergent functional networks related to antidepressant effects in major depressive disorder (12), and brain networks associated with schizophrenia symptoms (13). However, research on the network localization of anhedonia remains limited.

Transcranial magnetic stimulation (TMS) has shown promise in targeting neural circuits, with evidence suggesting that interventions focused on symptom-related circuits may be more effective in improving specific symptoms (14, 15). Selecting TMS targets based on the anhedonia-related damage network holds significant potential, as it considers symptom-specific neural pathways. However, no studies have yet integrated anhedonia-related damage network with TMS.

In this study, we first identify the damage network associated with anhedonia by mapping common functional impairments reported in neuroimaging studies, creating a convergent model of the anhedonia-related damage network through FCNM. More importantly, we explore how anhedonia-related damage network applied in the TMS treatment.

## 2. Materials and Methods

### 2.1. Study Search, Selection, and Extraction

The study search, selection, and extraction processes followed the consensus guidelines (16) and the Preferred Reporting Items for Systematic Reviews and Meta-Analyses (PRISMA) statement (17). This protocol has been registered with PROSPERO (registration number: CRD42022361560).

We performed a comprehensive and systematic literature search across PubMed, Web of Science, and Scopus to identify studies on FC associated with anhedonia. The search, which included studies published before August 29, 2022, used the keywords “anhedoni*”, “rest*” and “connect*”. We included studies that investigated anhedonia-related FC using seed-to-whole-brain voxel designs, and only studies with the same seed reported in more than five articles were included in the analysis. Coordinates from clusters derived from seed-based voxel-level analyses associated with anhedonia were extracted. Further details on the search, selection, and extraction procedures are provided in the Supplementary Methods.

### 2.2. Healthy Individual Dataset and fMRI Data Acquisition and Preprocessing

We selected 1,000 healthy individuals from the Brain Genomics Superstruct Project (https://dataverse.harvard.edu/dataverse/GSP), all of whom had resting-state fMRI scans and completed demographic and health questionnaires. Detailed demographic information is provided in Supplementary Table S1, and MRI scanning parameters are also outlined in Supplementary Methods.

Resting-state fMRI data were preprocessed using DPARSF software (http://rfmri.org/DPARSF), following the standard preprocessing pipeline recommended by the software. The first 10 volumes of each participant’s fMRI scan were discarded, and slice timing correction was applied. Realignment was performed to correct for head movement. T1-weighted structural images were co-registered with functional images, followed by segmentation and normalization using the DARTEL method. Nuisance covariates, including white matter (WM) and cerebrospinal fluid (CSF) signals, global signal, head motion, and polynomial trends, were regressed out. Head motion was corrected using Friston’s 24-parameter model. Finally, the datasets were band-pass filtered within the frequency range of 0.01 to 0.1 Hz.

### 2.3. Functional Connectivity Network Mapping

The FCNM method (9) was employed to identify the convergent neural circuits associated with anhedonia. First, 4-mm spheres were created around the peak coordinates derived from seed-based whole-brain voxel-level analyses related to anhedonia in each experiment and were then combined to form a combined seed. FC maps for the combined seed were computed for each individual, and these subject-level FC maps were compared to zero using a one-sample t-test with a threshold of t > 3 (11). Next, the thresholded t-maps were binarized, and the resulting binarized maps were combined to generate probability maps of the anhedonia-related FCNM, which were then thresholded at 60%, that is anhedonia-related damage network.

### 2.4. Subgroup Analysis

To determine whether the anhedonia-related damage network in major depressive disorder (MDD) aligns with the anhedonia-related damage network identified in the main analysis, we conducted an FCNM analysis exclusively on the MDD studies included in this study.

### 2.5. Transcriptomic Analysis

We explored the relationship between the anhedonia-related FCNM and gene expression to gain insights into the molecular mechanisms underlying the anhedonia-related damage network. Gene expression data were obtained from the Allen Human Brain Atlas (AHBA) dataset (http://human.brain-map.org), which includes over 20,000 gene expression measurements derived from 3,702 tissue samples (18). These samples were collected from six human donor brains, aged 24 to 57 years, with no reported neuropsychiatric or neuropathological history.

The microarray expression data were preprocessed using the abagen toolbox (19), which offers reproducible workflows for gene expression data processing following established guidelines. Standardized preprocessing workflows were applied. The preprocessing steps included: 1) intensity-based filtering to exclude probes that did not exceed background noise in more than 50% of the samples; 2) selection of a representative probe for each gene based on the most consistent regional variation across the six donor brains, as quantified by Differential Stability; 3) extraction of microarray samples from gray matter; and 4) gene normalization for all samples within structural classes, with aggregation conducted for each brain region, first independently for each donor and then across donors (19). This process resulted in gene expression data for 15,609 genes across 3,340 regions.

We applied partial least squares (PLS) regression in MATLAB to assess whether gene expression can predict the anhedonia-related FCNM (20). We first extracted the unthresholded probability maps of the anhedonia-related FCNM for the 3,340 gene sample coordinates, using a 4 mm radius. Our analysis included 3,340 brain regions, with 15,609 gene expression values as predictors (X) and the corresponding anhedonia-related FCNM scores for these 3,340 regions as the response variable (Y). The fitted PLS model generated several components, each of which represents a linear combination of the predictor variables explaining variance in the response variable. We retained the component that explained the most variance in the response variable. The significance of the explained variance was assessed via permutation tests, with spatial autocorrelation correction applied. To further refine the model, we used bootstrapping to assess and correct errors in estimating each gene’s contribution to the PLS component. Z-scores for each gene’s weight in the PLS model were calculated by the ratio of raw weight to bootstrapping error, and genes were ranked based on their contribution.

Functional enrichment analysis was performed to explore the functional annotations of genes associated with the anhedonia-related FCNM. This analysis was conducted using FUMA (https://fuma.ctglab.nl/). Significant genes (P_Bonferroni_ < 0.05) were submitted to FUMA for Gene Ontology (GO) enrichment analysis. To control for multiple comparisons, we applied the Benjamini-Hochberg False Discovery Rate (BH-FDR) method, setting the threshold for significant functional annotations at P < 0.05.

### 2.6. Anhedonia-Related Damage Network Applied in TMS Treatment

We used TMS treatment dataset to explore and expand the clinical application of the anhedonia-related damage network.

#### 2.6.1 TMS Treatment Dataset

A total of 35 adult participants were recruited from Tianjin Anding Hospital between November 2022 and December 2024 (clinical trial number: ChiCTR2100054793). All participants provided written informed consent, and the study was approved by the Ethics Committee of Tianjin Anding Hospital. Demographic and clinical information are provided in Supplementary Table S1.

Participants had a baseline 17-item Hamilton Rating Scale for Depression (HAMD-17) scores ≥ 12 and completed 20 sessions of Intermittent Theta-Burst Stimulation (iTBS), a form of high-frequency TMS with short bursts of stimulation, with 2 sessions per day over 10 consecutive days. Each session consisted of 3,000 pulses. The stimulation target was the left dorsolateral prefrontal cortex (DLPFC). Anhedonia was measured using the Snaith-Hamilton Pleasure Scale (SHAPS). The inclusion and exclusion criteria, data collection procedures, MRI parameters, and treatment details are provided in the Supplementary Methods. FMRI preprocessing followed the same procedures as those used in the GSP dataset.

#### 2.6.2 Baseline Associations Among Mean NAc FC in MDD, Anhedonia-Related FCNM, and Anhedonia Scores

We first examined the baseline association between seed-based FC in MDD patients, anhedonia-related FCNM, and anhedonia scores. Since only studies using the nucleus accumbens (NAc) as the seed met the eligibility criteria for FCNM analysis following a systematic search and selection, the anhedonia-related damage network was identified exclusively based on anhedonia-related NAc FC locations. Thus, we focused solely on NAc FC in the TMS dataset.

Based on the mean SHAPS scores, participants were categorized into high and low anhedonia groups. The characteristics, including age, gender, education, HAMD-17, and HAMA-14, showed no significant differences and were comparable between the two groups (Supplementary Table S2).

We calculated FC between NAc and whole-brain cortical regions in the TMS dataset. First, we extracted time-series data from the whole brain cortical regions according to the Schaefer atlas, along with data from the NAc (MNI coordinates: ± 9, 9, -8 with a 4 mm radius, which are commonly used in studies included for FCNM analysis). Functional connectivity between the NAc and all cortical regions from the Schaefer atlas was calculated using Fisher’s z-transformed Pearson correlation coefficients, yielding a 1 × 400 matrix representing the functional connectivity between the NAc and the whole brain cortical regions. We also mapped the anhedonia-related FCNM onto the Schaefer atlas, providing an anhedonia FCNM score for each of the 400 whole brain cortical regions.

We then examined the relationship between mean NAc FC and the FCNM score within each group using Spearman correlation. Additionally, we compared the mean NAc FC in the anhedonia-related damage network between the high and low anhedonia groups using the Mann-Whitney U test.

#### 2.6.3 Changes in Mean NAc FC Within the Anhedonia-Related Damage Network Following TMS Treatment and Its Association with Anhedonia Improvement

After excluding participants with SHAPS scores of 0 or 1, individuals were classified based on the percentage change in their SHAPS scores, calculated as (baseline - after scores) / baseline scores. A change percentage greater than 0.5 was used to differentiate between the response and non-response groups. The characteristics, including age, gender, education, HAMD-17, HAMA-14, and SHAPS, showed no significant differences and were comparable between the two groups (Supplementary Table S3). We then compared the change in mean NAc FC within the anhedonia-related damage network (after treatment mean NAc FC−bsaeline mean NAc FC) after TMS treatment between the response and non-response groups. Additionally, the change in mean NAc FC was correlated with improvements in efficacy for reducing anhedonia.

#### 2.6.4 Predicting TMS Treatment Efficacy Using Baseline Mean NAc FC Within the Anhedonia-Related Damage Network

We investigated whether baseline mean NAc FC within the anhedonia-related damage network could predict the efficacy of TMS treatment for anhedonia using a support vector machine (SVM) approach. Mean NAc FC was used to distinguish responders from non-responders, with model performance evaluated through 10-fold cross-validation. Additionally, we used the predictive performance of subjective SHAPS scores as a comparison, applying the same SVM model parameters.

## 3. Results

### 3.1. Included Studies

Out of an initial pool of 2,638 studies, seven studies (eight experiments) were selected for FCNM analysis. These studies focused on the NAc, with the majority examining individuals with MDD. Specifically, 4 studies focused solely on MDD, 1 on PTSD (Post-Traumatic Stress Disorder), 1 on PTSD + MDD, and 1 on low social anhedonia. In most studies (four studies), the NAc was defined using peak MNI coordinates (±9, 9, -8). One study adopted the NAc coordinates from Xia’s study (21), and two studies used the bilateral NAc from the Harvard-Oxford subcortical atlas. The study selection process is illustrated in Supplementary Figure S1, with further details provided in Supplementary Table S4.

### 3.2. Anhedonia-Related Damage Network

The anhedonia-related damage network encompasses a wide range of brain regions, primarily including the superior frontal cortex, caudate, cingulate gyrus, angular gyrus, cerebellum Crus II, mid temporal cortex, and the frontal orbital cortex (Figure 1). In terms of brain networks, the anhedonia-related damage network was predominantly linked to the default mode network (DMN, r = 0.302) and the reward network (r = 0.151), as determined using the decoder function of Neurosynth (https://neurosynth.org/). Furthermore, subgroup analysis in MDD revealed patterns strikingly similar to those observed in the main analysis (Supplementary Figure S2).

**Figure 1.**
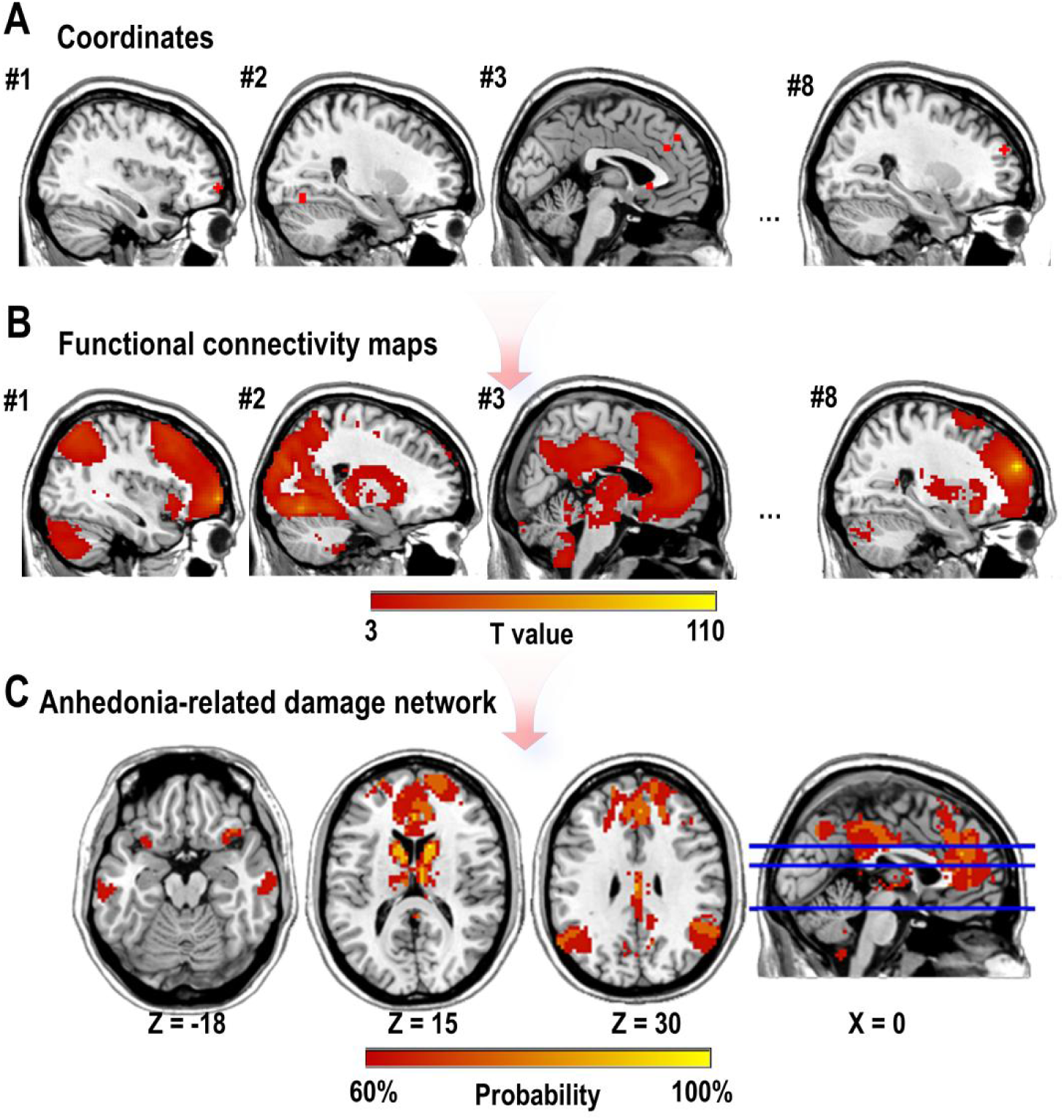
Anhedonia-Related Damage Network. A. Four-millimeter spheres were placed at abnormal coordinates derived from NAc-based whole-brain voxel-level analyses associated with anhedonia and then combined to form the combined seed. B. Brain regions significantly connected to the combined seed were identified using a large dataset of healthy individuals (n = 1000). FC maps for the combined seed were computed for each individual, and these subject-level FC maps were compared to zero using a one-sample t-test with a threshold of t > 3. C. The thresholded t-maps (t > 3) were binarized, and the resulting binarized maps were overlapped to create probability maps of the anhedonia-related damage network, which were thresholded at 60%.

### 3.3. Transcriptomic Analysis

We conducted transcriptomic analysis to identify genes associated with the anhedonia-related FCNM. The first component (PLS1) explained the most variance in the anhedonia-related FCNM, accounting for 10.7% of the variance, significantly more than expected by chance (Figure 2A, P_permutation_ < 0.001). The spatial distribution of PLS1 showed a positive correlation with the anhedonia-related FCNM (Figure 2B, r = 0.31, P < 0.001).

**Figure 2.**
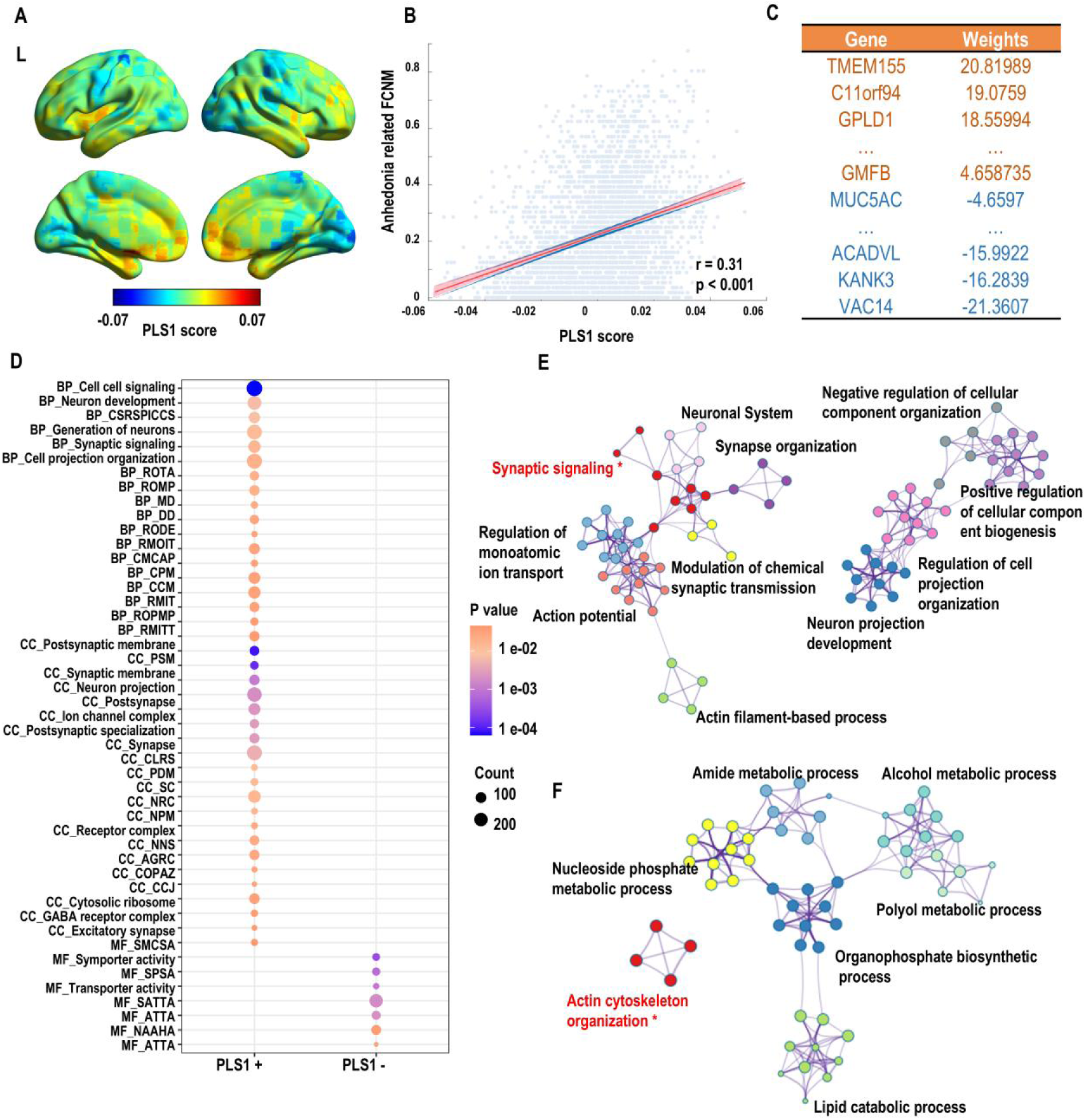
Neuroimaging Transcriptomics Analysis and Enrichment Analyses. A. The first component of Partial Least Squares (PLS1) scores. B. The significant spatial correlation between the PLS1 scores and the anhedonia-related functional connectivity network mapping (FCNM). C. The ranked gene weights of PLS1 (P_Bonferroni_ < 0.05) show that the gene weights of PLS1 are positive in orange and negative in blue. D. Significantly enriched Gene Ontology (GO) terms (P_Benjamini–Hochberg-FDR_ < 0.05) for the PLS1+ and PLS1-gene sets. The color of each circle represents the P value after Benjamini-Hochberg FDR correction, while the size of the circle indicates the number of genes enriched in the respective GO term. E. Metascape enrichment network for PLS1+. F. Metascape enrichment network for PLS1-. The Metascape enrichment network shows both intra-cluster and inter-cluster similarities of enriched annotations. Each term is represented by a node. Circular nodes represent enriched biological processes, with the size of the node reflecting the proportion of genes associated with the enriched process among all input genes. Nodes of the same color belong to the same cluster, and biological processes with a similarity greater than 0.3 are connected by edges. The most significant biological processes — related to synaptic signaling and actin cytoskeleton organization — are highlighted in red and marked with an asterisk (*), as they show the most significant enrichment. Abbreviations: BP_CSRSPICCS, Cell Surface Receptor Signaling Pathway Involved in Cell Cell Signaling; BP_ROTA, Regulation of Transporter Activity; BP_ROMP, Regulation of Membrane Potential; BP_MD, Membrane Depolarization; BP_DD, Dendrite Development; BP_RODE, Regulation of Dendrite Extension; BP_RMOIT, Regulation of Monoatomic Ion Transport; BP_CMCAP, Cardiac Muscle Cell Action Potential; BP_CPM, Cell Part Morphogenesis; BP_CCM, Cellular Component Morphogenesis; BP_RMIT, Regulation of Metal Ion Transport; BP_ROPMP, Regulation of Postsynaptic Membrane Potential; BP_RMITT, Regulation of Monoatomic Ion Transmembrane Transport; CC_PSM, Postsynaptic Specialization Membrane; CC_CLRS, Cytosolic Large Ribosomal Subunit; CC_PDM, Postsynaptic Density Membrane; CC_SC, Somatodendritic Compartment; CC_NRC, Neurotransmitter Receptor Complex; CC_NPM, Neuron Projection Membrane; CC_NNS, Neuron to Neuron Synapse; CC_AGRC, AMPA Glutamate Receptor Complex; CC_COPAZ, Cytoskeleton of Presynaptic Active Zone; CC_CCJ, Cell Cell Junction; MF_SMCSA, Solute Monoatomic Cation Symporter Activity; MF_SPSA, Solute Proton Symporter Activity; MF_SATTA, Secondary Active Transmembrane Transporter Activity; MF_ATTA, Amide Transmembrane Transporter Activity; MF_NAAHA, N Acylsphingosine Amidohydrolase Activity; MF_ATTA, Active Transmembrane Transporter Activity; NAc, nucleus accumbens; PLS, Partial Least Squares.

We identified 2,678 positively weighted genes (PLS1+ gene set) and 2,550 negatively weighted genes (PLS1-gene set), with both sets showing statistical significance (Figure 2C, P_Bonferroni_ < 0.05). The PLS1+ gene set was primarily enriched in processes related to synaptic signaling, neuron development, and ion transport. Specifically, Gene Ontology Biological Processes (GOBP) showed involvement in neuron development, synaptic signaling, cell signaling, and ion transport, while GO Cellular Components (GOCC) were mainly related to synaptic structures, postsynaptic membranes, and neurotransmitter receptor complexes. The PLS1-gene set was predominantly enriched in actin cytoskeleton organization (Figure 2D-E).

### 3.4. Anhedonia-Related Damage Network Applied in TMS Treatment

#### 3.4.1 Baseline Associations Among Mean NAc FC in MDD, Anhedonia-Related FCNM, and Anhedonia Scores

We found that the baseline mean NAc FC in the high-level anhedonia group was negatively correlated with the anhedonia FCNM scores, while in the low-level anhedonia group, the correlation was positive (Figure 3A). This suggests that a higher probability of anhedonia-related damage network is associated with more negative baseline NAc FC in the high-level anhedonia group and more positive NAc FC in the low-level anhedonia group. Individuals with high-level anhedonia exhibited lower baseline mean NAc FC within the anhedonia-related damage network compared to the low-level anhedonia group (Figure 3B).

**Figure 3.**
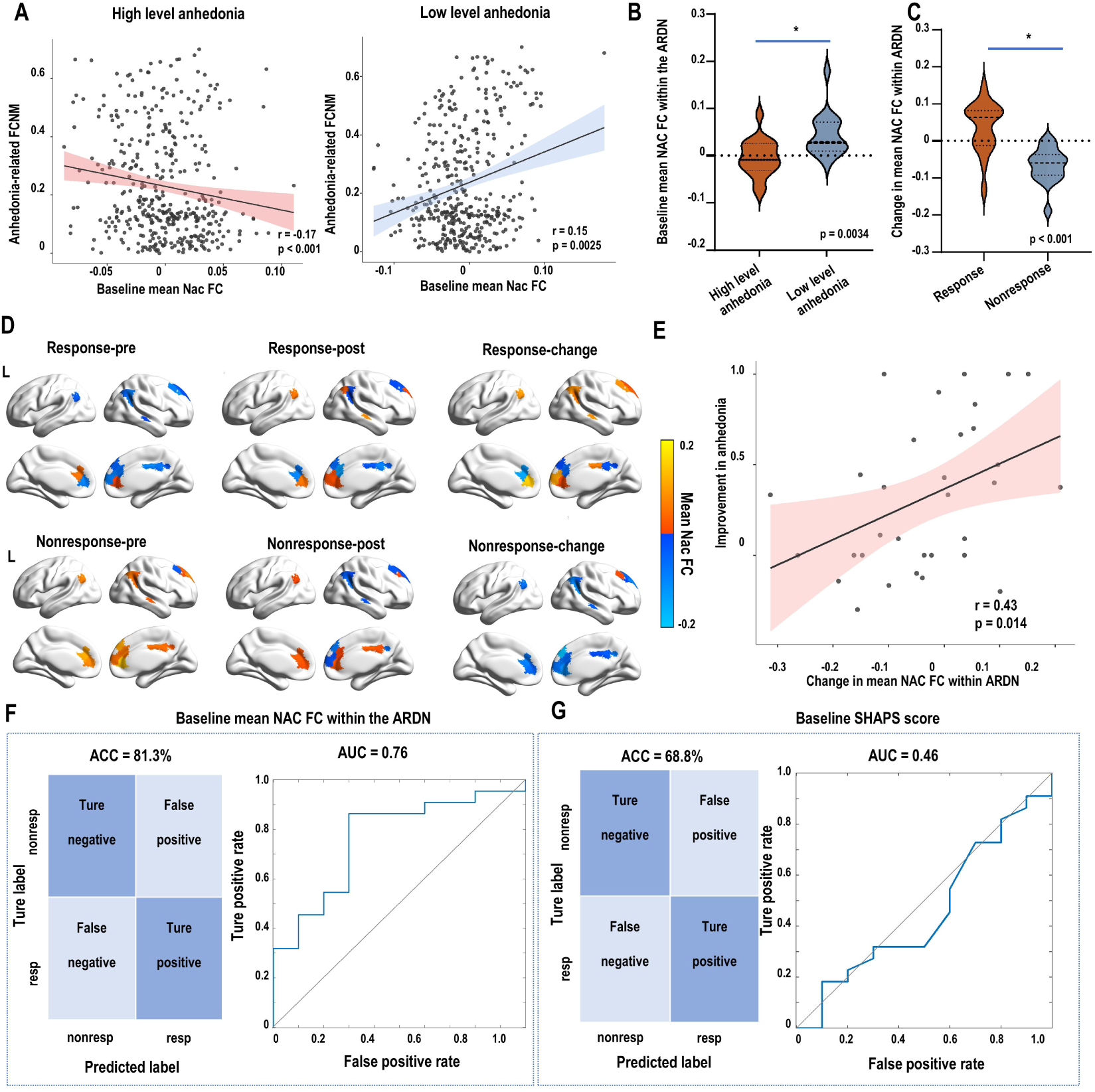
Application of the Anhedonia-Related Damage Network in TMS Treatment. A. The baseline mean NAc FC in the high-level anhedonia group was negatively correlated with the anhedonia FCNM scores, while in the low-level anhedonia group, the correlation was positive. This suggests that a higher probability of anhedonia-related damage network is associated with more negative baseline NAc FC in the high-level anhedonia group and more positive NAc FC in the low-level anhedonia group. B. Individuals with high-level anhedonia exhibited lower baseline mean NAc FC within the anhedonia-related damage network compared to the low-level anhedonia group. C. Changes in mean NAc FC within the anhedonia-related damage network varied between the treatment response and non-response groups. D. In the response group, changes in mean NAc FC within the anhedonia-related damage network shifted from a more negative pattern to a more positive one, whereas in the non-response group, the opposite pattern was observed. E. Changes in mean NAc FC within the anhedonia-related damage network were positively correlated with improvements in anhedonia symptoms. F. The baseline mean NAc FC within the anhedonia-related damage network was also found to effectively classify anhedonia responders from non-responders. The left panel shows the confusion matrix for the classifier, while the right panel presents the receiver operating characteristic (ROC) curve, demonstrating that the mean NAc FC within the anhedonia-related damage network performed significantly better than subjective baseline SHAPS scores (Figure 3G) for predicting treatment outcomes. Abbreviation: FCNM, functional connectivity network mapping; NAc, nucleus accumbens; FC, functional connectivity; ARDN, anhedonia-related damage network; pre, pre-TMS treatment; post, post-TMS treatment; resp, response, nonresp, nonresponse; ACC, accuracy; AUC, area under the receiver operating characteristic curve.

#### 3.4.2 Changes in Mean NAc FC Within the Anhedonia-Related Damage Network Following TMS Treatment and Its Association with Anhedonia Improvement

The response group showed an increase in the mean NAc FC within the anhedonia-related damage network following TMS treatment, whereas the non-response group showed a decrease (Figure 3C-D). Moreover, the increase in mean NAc FC within the anhedonia-related damage network was significantly correlated with improved efficacy in the treatment of anhedonia (Figure 3E).

#### 3.4.3 Predicting TMS Treatment Efficacy Using Baseline Mean NAc FC Within the Anhedonia-Related Damage Network

We found that baseline mean NAc FC within the anhedonia-related damage network can predict the efficacy of TMS treatment for anhedonia. A classifier using functional connectivity to distinguish responders from non-responders achieved an accuracy of 81.3% and an AUC of 0.76, evaluated through 10-fold cross-validation. In comparison, the subjective SHAPS scores performed poorly with an accuracy of 68.8% and an AUC of 0.46, using the same model parameters.

## 4. Discussion

This study identified an anhedonia-related damage network, primarily between the NAc and DMN. Transcriptomic analysis revealed enrichment in genes linked to synaptic signaling, neuron development, ion transport, and actin cytoskeleton. We found that individuals with high-level anhedonia exhibited lower baseline mean NAc FC within the anhedonia-related damage network compared to the low-level anhedonia group. TMS treatment increased mean NAc FC within the anhedonia-related damage network in the response group, which correlated with better outcomes in anhedonia. Baseline NAc FC also accurately predicted TMS response, outperforming subjective SHAPS scores. These results highlight the critical role of the anhedonia-related damage network in anhedonia patients, as well as its underlying mechanisms and predictive value for TMS in treating anhedonia.

This study identified an anhedonia-related damage network, primarily between the NAc and DMN, suggesting that dysfunction in the connection between the NAc and DMN plays a key role in anhedonia. This finding is highly consistent with previous research. Some studies have shown that the severity of anhedonia is correlated with the FC between the NAc and key DMN hubs (8, 22, 23). A study found that the severity of anhedonia was associated with the FC between the NAc and both the anterior cingulate cortex and the precuneus (8). Additionally, another study on subthreshold depression reported a positive association between increased FC of the DMN and NAc and a loss of interest (23). Furthermore, reduced connectivity between the NAc and the DMN has also been linked to reward deficits (24).

The gene set associated with anhedonia is primarily enriched in synaptic signaling, neuron development, ion channels, and actin cytoskeleton organization. Synaptic signaling and neuron development are fundamental to FC, and their dysfunction has been widely implicated in psychiatric conditions, including anhedonia (25, 26). Furthermore, ketamine, a fast-acting treatment for anhedonia, promotes synaptogenesis and reverses synaptic deficits caused by chronic stress, emphasizing the importance of synaptic dysfunction as a therapeutic target (27). The actin cytoskeleton plays a pivotal role in synaptic plasticity by enabling dynamic changes in synapse morphology in response to neuronal activity, which is crucial for synaptic transmission and plasticity (28). Ion channels also play a critical role in various neuronal activities, including growth, differentiation, signal transduction, and neurotransmitter release (29). Modulation of potassium channels, such as KCNQ, has been associated with restoring dopamine signaling and shows promise as an antianhedonic treatment for anhedonia (30).

We found that MDD patients with high anhedonia exhibited negative FC between the NAc and the anhedonia-related damage network, primarily within the DMN. Furthermore, we found that TMS treatment could enhance the reduced NAc-DMN connectivity, and this improvement was associated with better treatment efficacy for anhedonia. These findings are consistent with previous studies showing that high-frequency repetitive TMS increases FC between the NAc and various brain regions, including the fusiform gyrus, inferior temporal gyrus, calcarine fissure, superior occipital gyrus, and lingual gyrus, which are also primarily part of the DMN (31). A report found that NAc FC with a DMN node (the precuneus) was correlated with anhedonia improvement in escitalopram non-responders who received adjunctive aripiprazole (32). Additionally, a study proposed that an increase in NAc-dorsal anterior cingulate cortex connectivity from pre-treatment to post-treatment could distinguish responders from non-responders (33).

In our study, the anhedonia-related damage network not only provides insights into the mechanisms underlying anhedonia but also suggests that baseline mean NAc functional connectivity within this network may have the potential to partially predict treatment efficacy. Specifically, we found that baseline NAc functional connectivity within this network could be a potential predictor of treatment response, which may have practical implications for clinical decision-making and support more personalized treatment strategies for patients with anhedonia.

However, there are several limitations to consider. First, this study did not explore TMS targets based on the anhedonia-related damage network, which holds significant promise as it accounts for symptom-specific neural pathways. Since our study targeted only the DLPFC, which is not part of the anhedonia-related damage network, we were unable to examine a crucial aspect-whether the anhedonia-related damage network could guide TMS target selection and improve treatment efficacy. This remains a highly promising direction, and we hope future studies will investigate it further. Second, the sample size for the studies included in our analysis was limited. Similarly, the TMS dataset sample was also small, and future studies with larger samples are needed to validate these findings. Third, the gene expression data were derived from six healthy adult donors without MDD, which could impact the interpretation of the connectome-transcriptome associations in the context of depression.

## 5. Conclusions

This study identified an anhedonia-related damage network, primarily characterized by disrupted connectivity between the NAc and DMN. Our findings indicate that individuals with high-level anhedonia exhibit lower baseline NAc functional connectivity within this network. Notably, TMS treatment significantly increased NAc functional connectivity within the anhedonia-related damage network in responders, with these changes correlating with greater improvements in anhedonia symptoms. Furthermore, baseline NAc functional connectivity emerged as a potential predictor of TMS treatment response. These results underscore the critical role of the anhedonia-related damage network in the neuropathology of anhedonia and its predictive value for TMS efficacy. Future studies should further investigate the therapeutic implications of this network-based approach to optimize TMS target selection and improve treatment outcomes.

## Acknowledgements

Not applicable.

