## Supplemental file for "Mapping Anhedonia-Related Damage Network: Insights for TMS Treatment"

*Supplemental Information*

**Summary**

**Supplementary Methods**

***Search, selection, and extraction***
***MRI scanning parameters of the Brain Genomics Superstruct Project (GSP)***
***The inclusion and exclusion criteria, data collection procedures, MRI parameters, and treatment details of TMS treatment dataset***

***Supplementary Figure S1. Study selection flowchart.***
***Supplementary Figure S2. The Anhedonia-Related Damage Network of Major Depressive Disorder***

***Supplementary Table S1. Demographic and clinical information from the Brain Genomics Superstruct Project dataset and the TMS treatment dataset.***
***Supplementary Table S2. Demographic and clinical information for the high level anhedonia and low level anhedonia groups in the TMS treatment dataset.***
***Supplementary Table S3. Demographic and clinical information for the response and nonresponse groups in the TMS treatment dataset.***
***Supplementary Table S4. Characteristics and Coordinates of the Studies Included.***

**Supplementary methods**

***Search, selection, and extraction***

The study search, selection, and extraction processes followed the consensus guidelines (1) and the Preferred Reporting Items for Systematic Reviews and Meta-Analyses (PRISMA) statement (2). The protocol has been registered with PROSPERO (registration number: CRD42022361560).

We conducted a comprehensive and systematic literature search across PubMed, Web of Science, and Scopus to identify studies on functional connectivity (FC) associated with anhedonia. The search, which included studies published before August 29, 2022, used the keywords "anhedoni*," "rest*," and "connect*." The search was not limited to any specific diagnosis, as we assumed that a common network related to anhedonia exists across different diseases.

Eligible studies had to meet the inclusion criteria, which were first assessed through preliminary screening based on the title and abstract. Studies were included if they reported anhedonia-associated FC, were resting-state fMRI studies published in English or Chinese, and were case-control studies, cohort studies, or randomized controlled trials (RCTs) that allowed extraction of original neuroimaging data. Exclusion criteria at this stage included studies focusing solely on structural results, unpublished studies, case reports, conference abstracts, narrative or systematic reviews, meta-analyses, letters, and other secondhand sources. After preliminary screening, a second phase of full-text screening was performed. Studies were included if they used seed-to-whole-brain voxel designs and reported coordinates in MNI space or Talairach space, and only those with the same seed reported in more than five articles were considered. Studies were excluded if they involved overlapping samples or participants with known severe neurological conditions or head trauma.

The search, selection, and extraction processes were conducted independently by Xuezhen Zhang and Xinyan Rong, with disagreements resolved by a third reviewer (Chengfeng Chen). A flow diagram of the study selection process is presented in Supplementary Figure S1. We also extracted author information, seed definitions, and coordinates from clusters derived from seed-based voxel-level analyses associated with anhedonia, as well as information on population characteristics and sample sizes.

***fMRI scanning parameters of GSP***

We selected 1,000 healthy individuals from the Brain Genomics Superstruct Project (https://dataverse.harvard.edu/dataverse/GSP). We utilized structural MRI scans and the second set of resting-state fMRI scans from the GSP dataset, which were acquired on matched 3T Tim Trio scanners (Siemens, Erlangen, Germany) with a 12-channel phased-array head coil.

The structural MRI scans were acquired using a T1-weighted Multi-Echo Magnetization Prepared Rapid Gradient Echo (MEMPRAGE) sequence with the following parameters: a repetition time (TR) of 2.2 seconds, echo times (TE) of 1.5 ms for the first image, and 7.0 ms for the fourth image, a flip angle of 7°, an inversion time (TI) of 1.1 seconds, a slice thickness of 1.2 mm, 144 slices, and an isotropic resolution of 1.2 × 1.2 × 1.2 mm.

The resting-state fMRI scans were acquired using a gradient-echo echo-planar imaging sequence, a TR of 3.0 seconds, a TE of 30 milliseconds, a flip angle of 85°, and a slice thickness of 3.0 mm. The fMRI data consisted of 47 slices with an isotropic resolution of 3.0 × 3.0 × 3.0 mm (3).

***The inclusion and exclusion criteria, data collection procedures, MRI parameters, and treatment details of TMS treatment dataset***

A total of 36 adult participants were recruited from Tianjin Anding Hospital between November 2022 and December 2024 (clinical trial number: ChiCTR2100054793). All participants provided written informed consent, and the study was approved by the Ethics Committee of Tianjin Anding Hospital. Demographic information is provided in Table S1 of the Supplementary Materials.

The inclusion criteria for this study included: 1) Aged between 18 and 60 years; 2) Diagnosed with major depressive disorder (MDD) by psychiatrists based on Diagnostic and Statistical Manual of Mental Disorders (Fifth Edition) (DSM-5), and this diagnosis was confirmed through a Mini International Neuropsychiatric Interview (MINI); 3) The 17-item Hamilton Depression Rating Scale (HAMD-17) scores of 12 or higher. 4) No adjustments to the medication type were made during the treatment period. The exclusion criteria for this study included: 1) Metal implants are placed above the chest, particularly in the intracranial or cardiac regions; 2) Presence of serious medical conditions, including neurological disorders, brain diseases, traumatic brain injury or surgery, epilepsy, or severe physical conditions such as malignant tumors, acute heart failure, multi-organ failure, and infectious diseases; 3) Presence of psychotic symptoms; 4) History of receiving repetitive transcranial magnetic stimulation (rTMS) or electroconvulsive therapy (ECT) within the last 6 months; 5) History of drug or alcohol abuse within the past year; 6) Inability to communicate, understand, or follow instructions normally, or inability to cooperate with treatment and evaluation; 7) Currently participating in other clinical trials involving drug or physical treatments.

Participants completed 20 sessions of intermittent Theta-Burst Stimulation (iTBS), a form of high-frequency TMS with short bursts of stimulation, with 2 sessions per day over 10 consecutive days. The TMS protocol used was: 50 Hz intraburst frequency, 5 Hz interburst frequency, 2-second stimulation, and 8-second rest. Each session consisted of 3,000 pulses (lasting 16 minutes and 12 seconds). The interval between the two sessions was no less than 50 minutes. The stimulation target was the left dorsolateral prefrontal cortex (DLPFC). Based on the standard "5 cm" method (MNI coordinates were -41, 16, 45) and targeting the dorsolateral prefrontal cortex (DLPFC) with the most negative functional connectivity to the posterior cingulate cortex (pgACC, MNI coordinates were -10, 42, 6), patients were randomly assigned to two groups.

The demographic information, including gender, age, education, and medication use, was self-reported by patients and collected at baseline. In addition, ratings and MRI scans were administered at baseline and again at the end of 10 days, following the completion of the twenty iTBS treatment sessions. Patient's depression severity was assessed using the HAMD-17 and the Snaith-Hamilton Pleasure Scale (SHAPS) was used to measure anhedonia. The Snaith-Hamilton Pleasure Scale (SHAPS) consists of 14 items, each with four response options (ranging from 1 to 4 points). Responses of '1' and '2' ('strongly agree' and 'agree') are scored as 0, while responses of '3' and '4' ('disagree' and 'strongly disagree') are scored as 1. The total score ranges from 0 to 14 points, with a higher score indicating a more severe level of anhedonia.

MRI data were acquired using Siemens Magnetom Prisma 3.0 T MRI scanners. For the 3D T1-weighted structural images, a Magnetization-Prepared Rapid Gradient Echo (MP-RAGE) sequence was employed with a repetition time (TR) of 2000 ms, echo time (TE) of 2.32 ms, field of view (FOV) of 230 × 230 $mm^2$, a matrix of 256 × 256, a flip angle of 8°, slice thickness of 0.90 mm, 208 slices, and a voxel size of 0.9 × 0.9 × 0.9 $mm^3$. For resting-state fMRI (rs-fMRI), a Gradient-Echo Echo-Planar Imaging (GE-EPI) sequence was used with a TR of 800 ms, TE of 30 ms, FOV of 208 × 208 $mm^2$, a matrix of 104 × 104, a flip angle of 56°, slice thickness of 2 mm, 72 slices, a voxel size of 2 × 2 × 2 $mm^3$, and 450 time points.

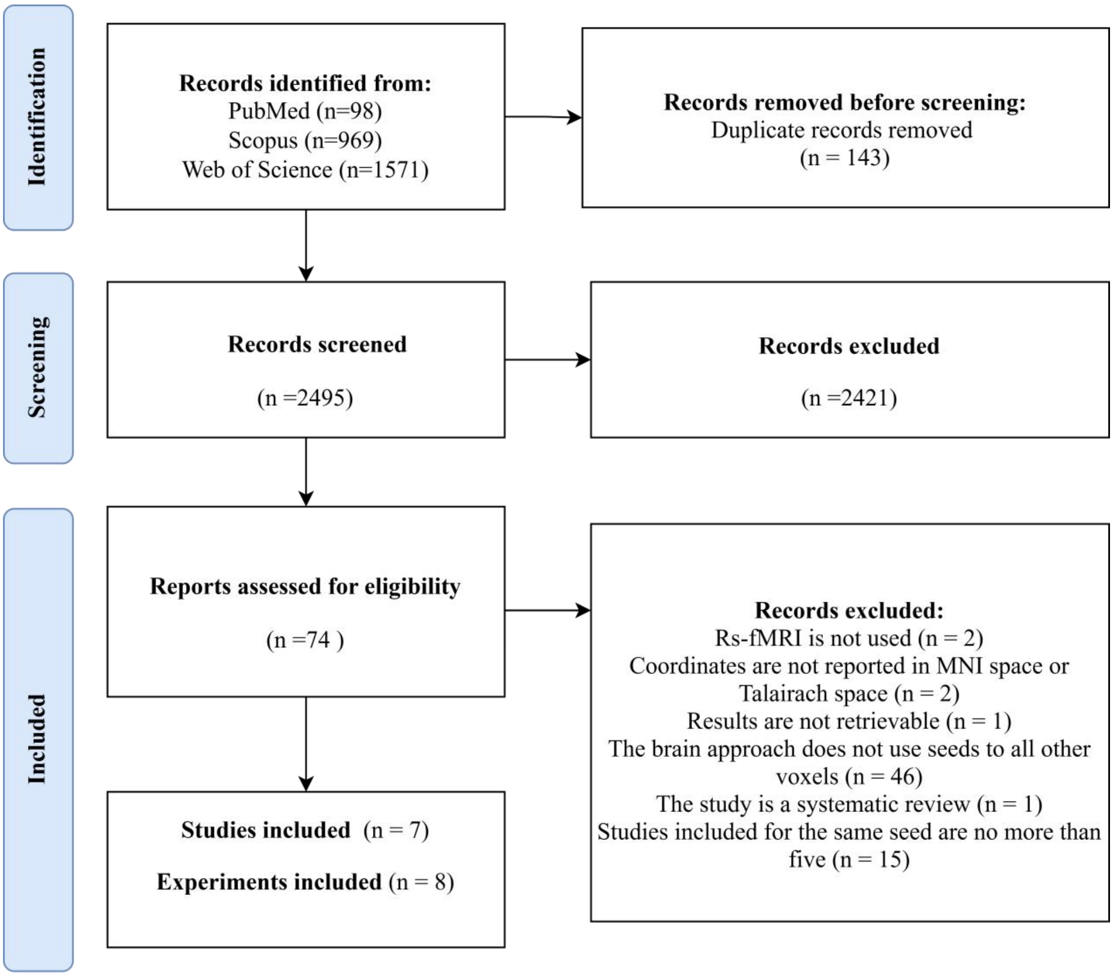


***Supplementary Figure S1. Study selection flowchart.***

Abbreviations: rs-fMRI: resting-state functional magnetic resonance imaging; MNI: Montreal Neurological Institute.

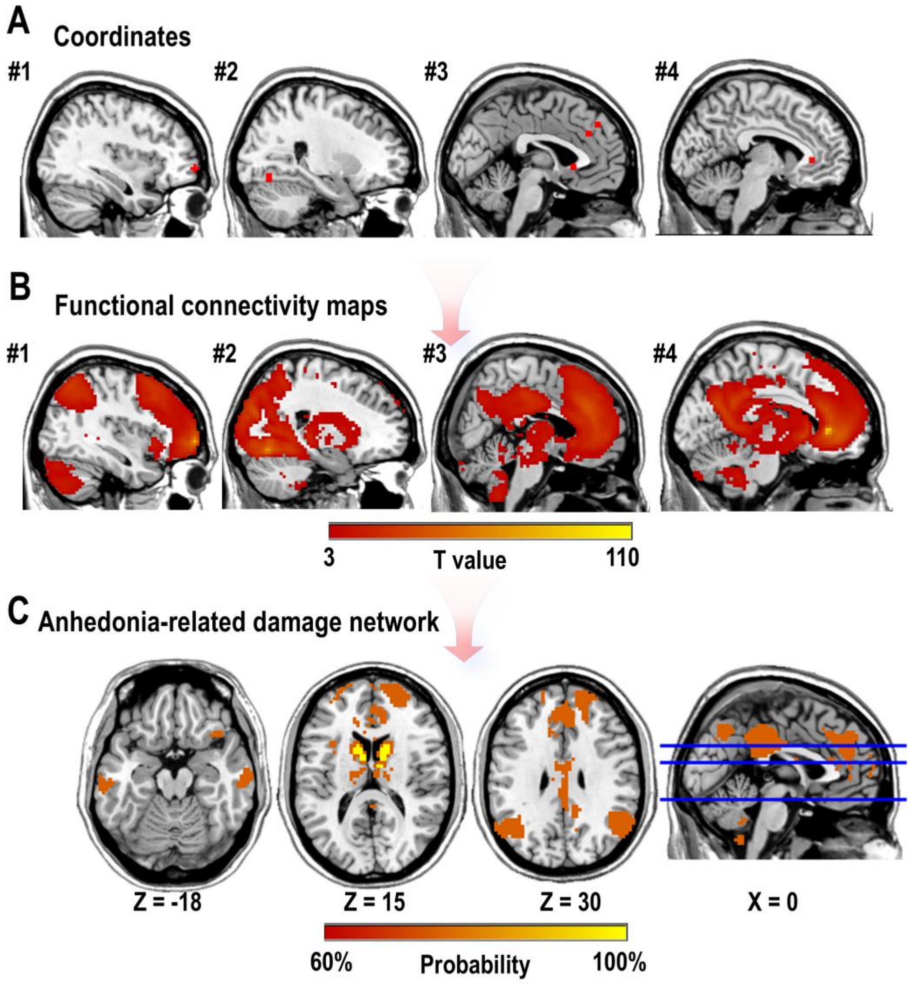


***Supplementary Figure S2. The Anhedonia-Related Damage Network of Major Depressive Disorder***

A. Four-millimeter spheres were placed at abnormal coordinates derived from NAc-based whole-brain voxel-level analyses associated with anhedonia in major depressive disorder and then combined to form the combined seed.

B. Brain regions significantly connected to the combined seed were identified using a large dataset of healthy individuals (n = 1000). Functional connectivity (FC) maps for the combined seed were computed for each individual, and these subject-level FC maps were compared to zero using a one-sample t-test with a threshold of t > 3.

C. The thresholded t-maps (t > 3) were binarized, and the resulting binarized maps were overlapped to create probability maps of the anhedonia-related damage network, which were thresholded at 60%.

***Supplementary Table S1. Demographic and clinical information from the Brain Genomics Superstruct Project dataset and the TMS treatment dataset.***

| **Dataset** | **Sample size** | **Age (years)** | **Sex (F/M)** | **Education (years)** | **Handedness (AMB/LFT/RHT)** | **HAMD-17** | **HAMA-14** |
|---|---|---|---|---|---|---|---|
| Brain Genomics Superstruct Project | 1000 | 21.5±2.85 | 580/420 | 14.5±1.86 | 7/77/916 | Na | Na |
| TMS treatment dataset | 35 | 34.8±12.0 | 21/14 | 21/14 | 0/0/35 | 20.9±4.91 | 21.8±7.60 |

Abbreviations: AMB, Ambidextrous; LFT, Left-handed; RHT, Right-handed; F, Female; M, Male; Na, Not applicable; HAMD-17, Hamilton Depression Rating Scale-17 items; HAMA-14, Hamilton Anxiety Rating Scale-14 items.

***Supplementary Table S2. Demographic and clinical information for the high level anhedonia and low level anhedonia groups in the TMS treatment dataset.***

| | High level anhedonia (n=21) | Low level anhedonia (n=14) | $Z/x^2$ | P-value |
|---|---|---|---|---|
| Age (years), mean ± SD | 34.3±11.9 | 35.6±12.6 | -0.253 | 0.801 |
| Sex (F/M) | 13/8 | 8/6 | 0.0794 | 0.778 |
| Education (years), mean ± SD | 14.2±3.46 | 14.4±4.27 | -0.203 | 0.839 |
| HAMD-17, mean ± SD | 21.9±4.44 | 19.5±5.39 | 1.34 | 0.181 |
| HAMA-14, mean ± SD | 22.0±6.04 | 21.4±9.73 | 0.590 | 0.555 |
| SHAPS, mean ± SD | 10.3±1.76 | 4.36±2.47 | 4.96 | <0.001 |

The characteristics, including age, education, HAMD-17, HAMA-14, and SHAPS, were analyzed using the Mann-Whitney U test due to non-normal distribution, and the Z statistic was used to present the results. Sex was analyzed using the Chi-square test.

Abbreviations: TMS, Transcranial Magnetic Stimulation; SD, Standard Deviation; F, Female; M, Male; HAMD-17, Hamilton Depression Rating Scale-17 items; HAMA-14, Hamilton Anxiety Rating Scale-14 items; SHAPS, Snaith-Hamilton Pleasure Scale

***Supplementary Table S3. Demographic and clinical information for the response and nonresponse groups in the TMS treatment dataset.***

| | Response (n=22) | Nonresponse (n=10) | $Z/x^2$ | P-value |
|---|---|---|---|---|

| | | | | |
|---|---|---|---|---|
| Age (years), mean ± SD | 34.8±10.7 | 34.9±13.1 | 0.264 | 0.792 |
| Sex (F/M) | 12/10 | 6/4 | 0.0831 | 0.773 |
| Education (years), mean ± SD | 13.9±4.01 | 14.2±3.91 | 0.306 | 0.760 |
| HAMD-17, mean ± SD | 23.0±4.94 | 20.3±4.54 | 1.43 | 0.152 |
| HAMA-14, mean ± SD | 22.6±7.28 | 21.3±7.65 | 0.550 | 0.582 |
| SHAPS, mean ± SD | 9.00±2.87 | 8.41±2.99 | 0.409 | 0.682 |

The response was defined as a greater than 50% change in SHAPS, while nonresponse was defined as less than a 50% change. We excluded cases where the baseline SHAPS score was 0 or 1.

The characteristics, including age, education, HAMD-17, HAMA-14, and SHAPS, were analyzed using the Mann-Whitney U test due to non-normal distribution, and the Z statistic was used to present the results. Sex was analyzed using the Chi-square test.

Abbreviations: TMS, Transcranial Magnetic Stimulation; SD, Standard Deviation; F, Female; M, Male; HAMD-17, Hamilton Depression Rating Scale-17 items; HAMA-14, Hamilton Anxiety Rating Scale-14 items; SHAPS, Snaith-Hamilton Pleasure Scale

***Supplementary Table S4. Characteristics and Coordinates of the Studies Included.***

| Experiments | Studies | X | Y | Z | Relationship with Anhedonia (Positive/Negative) | Seed | Seed Definition | Scale | Population | Sample |
|---|---|---|---|---|---|---|---|---|---|---|
| 1 | (Fan, Liu et al. 2021) (4) | 36 | 60 | -3 | N | NAc | Peak coordinated:±9，9，-8 | PAS | MDD | 66 |
| 2 | (Gabbay, Ely et al. 2013) (5) | -8 | 10 | 14 | N | NAc | Peak coordinated:±9，9，-9 | BDI-II (Items 4, 12) + CDRS-R (Item 2) | MDD | 21 |
| 2 | (Gabbay, Ely et al. 2013) (5) | 24 | -70 | -12 | N | NAc | Peak coordinated:±9，9，-10 | BDI-II (Items 4, 12) + CDRS-R (Item 3) | MDD | 21 |

| | | | | | | | | | | |
|---|---|---|---|---|---|---|---|---|---|---|
| 3 | (Gong, He et al. 2018) (6) | -27 | 54 | 16 | N | NAc | The bilateral NAc from Harvard-Oxford subcortical atlas | TEPS | MDD | 68 |
| 3 | (Gong, He et al. 2018) (6) | -3 | 44 | 41 | P | NAc | The bilateral NAc from Harvard-Oxford subcortical atlas | TEPS | MDD | 68 |
| 3 | (Gong, He et al. 2018) (6) | -3 | 35 | 31 | P | NAc | The bilateral NAc from Harvard-Oxford subcortical atlas | TEPS | MDD | 68 |
| 3 | (Gong, He et al. 2018) (6) | -32 | -74 | 49 | P | NAc | The bilateral NAc from Harvard-Oxford subcortical atlas | TEPS | MDD | 68 |
| 3 | (Gong, He et al. 2018) (6) | -41 | 9 | -2 | P | NAc | The bilateral NAc from Harvard-Oxford subcortical atlas | TEPS | MDD | 68 |
| 3 | (Gong, He et al. 2018) (6) | 43 | 15 | 1 | N | NAc | The bilateral NAc from Harvard-Oxford subcortical atlas | TEPS | MDD | 68 |
| 3 | (Gong, He et al. 2018) (6) | -17 | 35 | -12 | N | NAc | The bilateral NAc from Harvard-Oxford subcortical atlas | TEPS | MDD | 68 |
| 3 | (Gong, He et al. 2018) (6) | 11 | 35 | -11 | N | NAc | The bilateral NAc from Harvard-Oxford subcortical atlas | TEPS | MDD | 68 |

| | | | | | | | | | | |
|---|---|---|---|---|---|---|---|---|---|---|
| 3 | (Gong, He et al. 2018) (6) | -4 | 20 | -1 | N | NAc | The bilateral NAc from Harvard-Oxford subcortical atlas | TEPS | MDD | 68 |
| 3 | (Gong, He et al. 2018) (6) | 7 | 20 | -4 | N | NAc | The bilateral NAc from Harvard-Oxford subcortical atlas | TEPS | MDD | 68 |
| 4 | (Liu, Wang et al. 2021)-HC (7) | 4 | -58 | 18 | P | NAc | NAc from (Xia, Fan et al. 2017) (8) | TEPS | HC | 28 |
| 5 | (Liu, Wang et al. 2021)-MDD (7) | -6 | 34 | 2 | N | NAc | NAc from (Xia, Fan et al. 2017) (8) | TEPS | MDD | 23 |
| 6 | (Olson, Kaiser et al. 2018) (9) | 14 | 56 | 16 | P | NAc | The bilateral NAc from Harvard-Oxford subcortical atlas | SHAPS | PTSD | 51 |
| 7 | (Pessin, Philippi et al. 2021) (10) | -13 | -18 | 21 | N | NAc | Peak coordinated:±9，9，-10 | MASD-AD | PTSD | 71 |
| 8 | (Wang, Liu et al. 2016) (11) | 27 | 54 | 24 | P | NAc | Peak coordinated:±9，9，-10 | TEPS | Low socAnh | 30 |

Abbreviations: NAc, Nucleus Accumbens; N, Negative; P, Positive; PAS, Physical Anhedonia Scale; CDRS-R, Children's Depression Rating Scale–Revised; BDI-II, the Beck Depression Inventory, 2nd edition; TEPS, Temporal Experience of Pleasure Scale; SHAPS, the Snaith-Hamilton Pleasure Scale; MASD-AD, the Anhedonic Depression subscale of the Mood and Anxiety Symptoms Questionnaire; MDD, Major Depressive Disorder; HC, Healthy Control; PTSD, Post-Traumatic Stress Disorder; Low socAnh, Low Social Anhedonia.
